# Born with an abstract code for number: evidence from cross-modal and cross-format neural adaptation in the first three days of life

**DOI:** 10.64898/2026.05.08.723896

**Authors:** Marco Buiatti, Elena Eccher, Irene Petrizzo, Ugo Pradal, Fabrizio Taddei, Giorgio Vallortigara, Véronique Izard, Manuela Piazza

**Author notes:** Corresponding Author: Marco Buiatti.

## Abstract

One of the most astonishing abilities documented in human newborns is their capacity to spontaneously match sets of items on the basis of their number, even when they are presented in different modalities (auditory and visual) and formats (sequential and simultaneous). These behavioural findings suggest that the newborn brain comes to life endowed with a system that encodes number in an abstract, supramodal format, but direct neural evidence for such a representation is lacking. Using high-density EEG and a frequency-tagging paradigm, we measured neural entrainment to periodically presented visual arrays that were either numerically congruent or incongruent with previously familiarized and concurrently presented auditory sequences in 21 newborns (0–3 days old). The amplitude of neural entrainment to the visual arrays provided a robust index of cross-modal numerical congruency: it was significantly reduced for numerically congruent relative to incongruent stimuli. This cross-modal and cross-format numerosity-specific repetition suppression effect provides evidence for an abstract code for number in the newborn brain, reinforcing the view that number constitutes a foundational dimension of human perception since birth.

## Introduction

Human newborns, like other animals, enter the world equipped with a repertoire of instinctual abilities that rest on prior expectations about the structure of their environment^1,2^. Characterizing their nature and extent is critical because they provide the foundations that guide and constrain subsequent learning and cognitive development. Seminal work by Izard and colleaguereported that, already within the first few days of life, newborns hearing auditory sequences containing a fixed number of syllables looked longer at numerically congruent visual arrays of objects (simple geometrical shapes). These findings suggest an early sense of number that generalizes across sensory modalities and stimulus formats. However, the neural computations supporting this ability remain unknown. One possibility is that in the newborn brain numerosity is represented in an abstract, modality independent format, potentially by supramodal neurons that are equally activated regardless of whether a given numerosity is presented visually or auditorily, sequentially or simultaneously^5^. To probe the existence of such a neural code, we took advantage of the repetition-suppression phenomenon: When a specific feature of a stimulus is repeated, the responses of a neural population that encode that specific feature decrease, even in the context of other feature changes^6,7^. In light of this effect, the hypothesis of an abstract neural code for number implies a straightforward prediction: Repeated exposure to a given numerosity should induce neural repetition suppression even when numerical information is presented through different sensory modalities and formats. To test this prediction, we adapted the paradigm introduced by^3^ for a high-density EEG study in very young newborns (0–3 days old). Using a frequency-tagging approach^8^, we measured neural responses to visual arrays that were either numerically congruent or incongruent with previously familiarized and concurrently presented auditory sequences, thereby directly probing the neural basis of early numerical representations.

## Results

During an initial familiarization phase (1.5 min), 21 healthy newborns were exposed to auditory sequences of a fixed number of syllables (either 4 or 12). In the subsequent test phase, while auditory stimulation continued, visual arrays of 4 or 12 items were presented dynamically on the screen through a sinusoidal contrast modulation (0–100%) with a rate of 0.8 Hz (Fig. 1). Visual stimuli were delivered in alternating numerosity blocks, lasting up to 50 s, or until fixation was lost.

**Figure 1:**
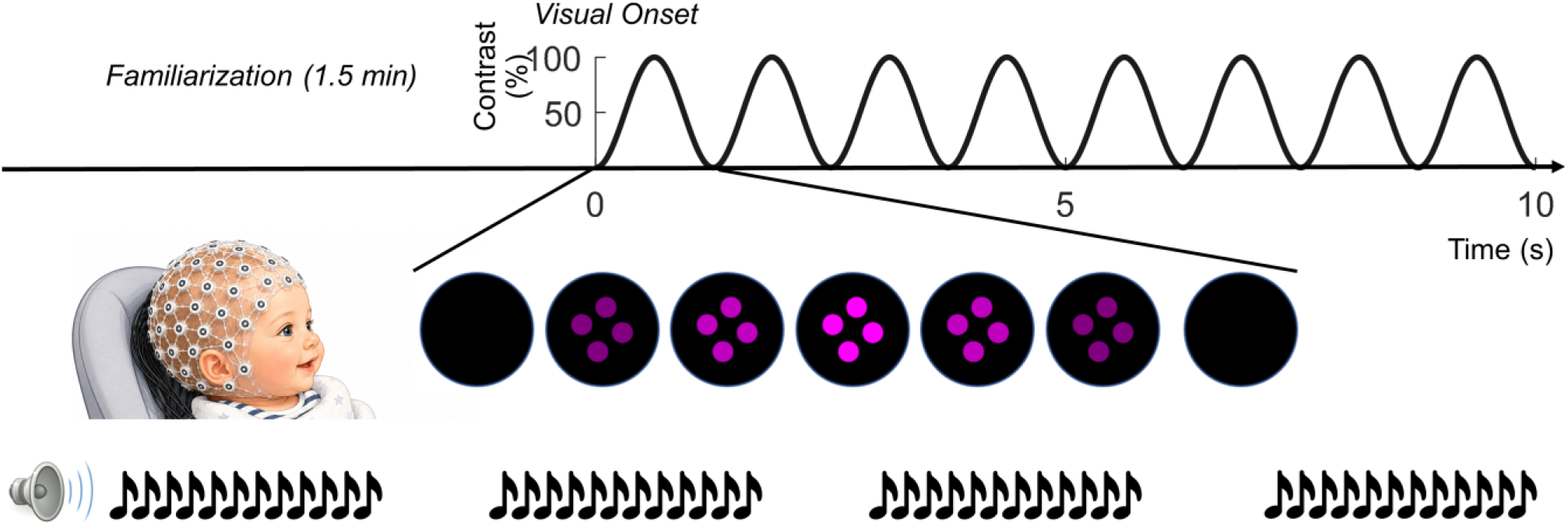
Experimental paradigm. During the first 1.5 minutes, newborns were familiarized with auditory sequences of a fixed number of syllables (4 or 12); they were then tested with visual arrays with the same or a different number of items (4 or 12) presented with sinusoidal contrast modulation (0 to 100%) at a rate of 0.8 Hz, while auditory sequences continued.

To assess the effect of cross-modal numerosity congruency, we quantified the response to the test visual stimuli using two complementary measures of entrainment (see Methods): (i) synchronization (phase-locking) to the stimulus oscillation, indexed by Inter-Trial Coherence (ITC), and (ii) amplitude of the entrained oscillations at the stimulus frequency, indexed by the Frequency-Tagged Response (FTR). Despite the short duration of the overall usable EEG data due to newborns’ limited visual attention (mean per subject: 135 s) and the presence of simultaneous, asynchronous auditory input, infants reliably engaged with the visual stimulation, as evident by both significant ITC and FTR at the stimulation frequency compared to background activity (both conditions merged, *P*_*corr*_ = 0.013 and *P*_*corr*_ = 0.003, respectively). The spatial distribution of these effects was highly consistent across the two measures, with overlapping posterior electrode clusters characteristic of visual cortical responses (Fig. 2).

**Figure 2:**
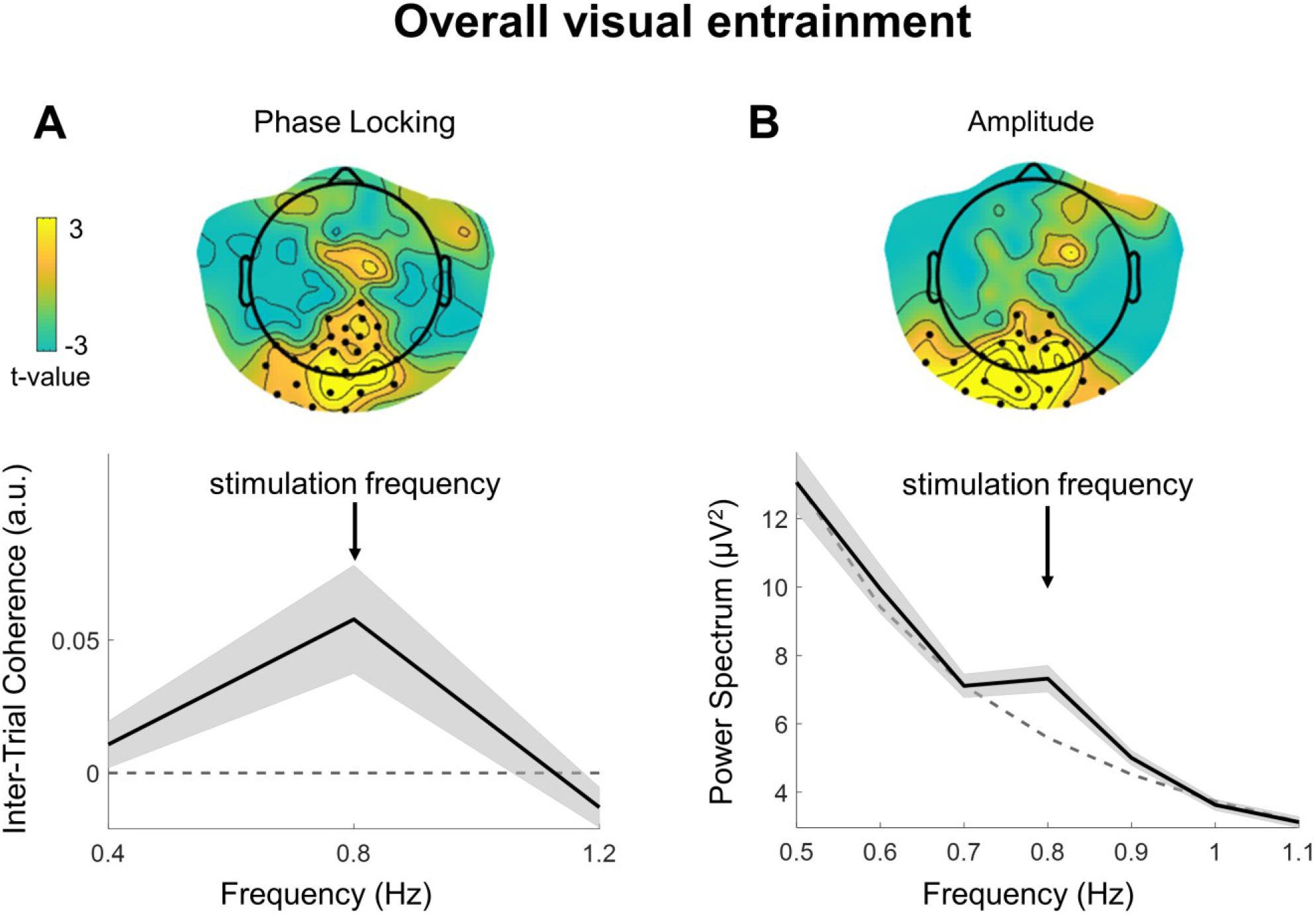
Overall visual entrainment. (A) Phase locking. Top: Statistical map (t-value, one-tailed t-test; black dots indicate statistically significant clusters (cluster-based permutation test, see Methods for details)) of the difference between ITC at the stimulation frequency (0.8 Hz) and the one computed from the null distribution generated from the same data. The map shows statistically significant phase locking (*P*_*corr*_ < 0.013) in a wide posterior area. Bottom: Spectral profile of the difference ITC averaged over the significant cluster (black line) ± SEM (gray shadow) across subjects: a clear peak above the null-distribution level (marked by the dashed line at zero) emerges at the stimulation frequency. (B) Amplitude of entrainment. Top: Statistical map (as above) of the difference between the Power Spectrum (PS) at the tag frequency (0.8 Hz) and the background power at the same frequency, significant (*P*_*corr*_ < 0.003) in a posterior cluster of electrodes overlapping with the one emerging from phase locking. Bottom: Mean of the single-subject difference between the PS and the background power averaged over electrodes belonging to the posterior cluster (black line) ± SEM (gray shadow), superposed on the average background power (dashed dark-gray line): a PS peak neatly emerges at the stimulation frequency, indicating significant entrainment.

We next investigated the effect of cross-modal numerical congruency on the test visual stimuli. No significant difference was observed in ITC between congruent and incongruent conditions (Fig. 3A, top panel; P>0.05 for all electrodes, no significant clusters detected), indicating that in both conditions the newborn brain temporally tracks the visual stimulation. In contrast, the FTR markedly differed, being significantly lower in the congruent condition compared to the incongruent one (Fig. 3B; left cluster: *P*_*corr*_ < 0.001, effect size *d =* 1.04; right cluster: *P*_*corr*_ < 0.014) in a large bilateral cluster of electrodes, compatible with an associative temporo-parietal source. This pattern is consistent with a cross-modal and cross-format numerosity-specific repetition suppression effect, whereby following a repeated presentation of a given numerosity, the neural populations encoding that numerosity exhibit a suppressed response to the numerically congruent stimuli, despite that such stimuli change in presentation modality (from auditory to visual) and format (from sequential to simultaneous).

**Figure 3:**
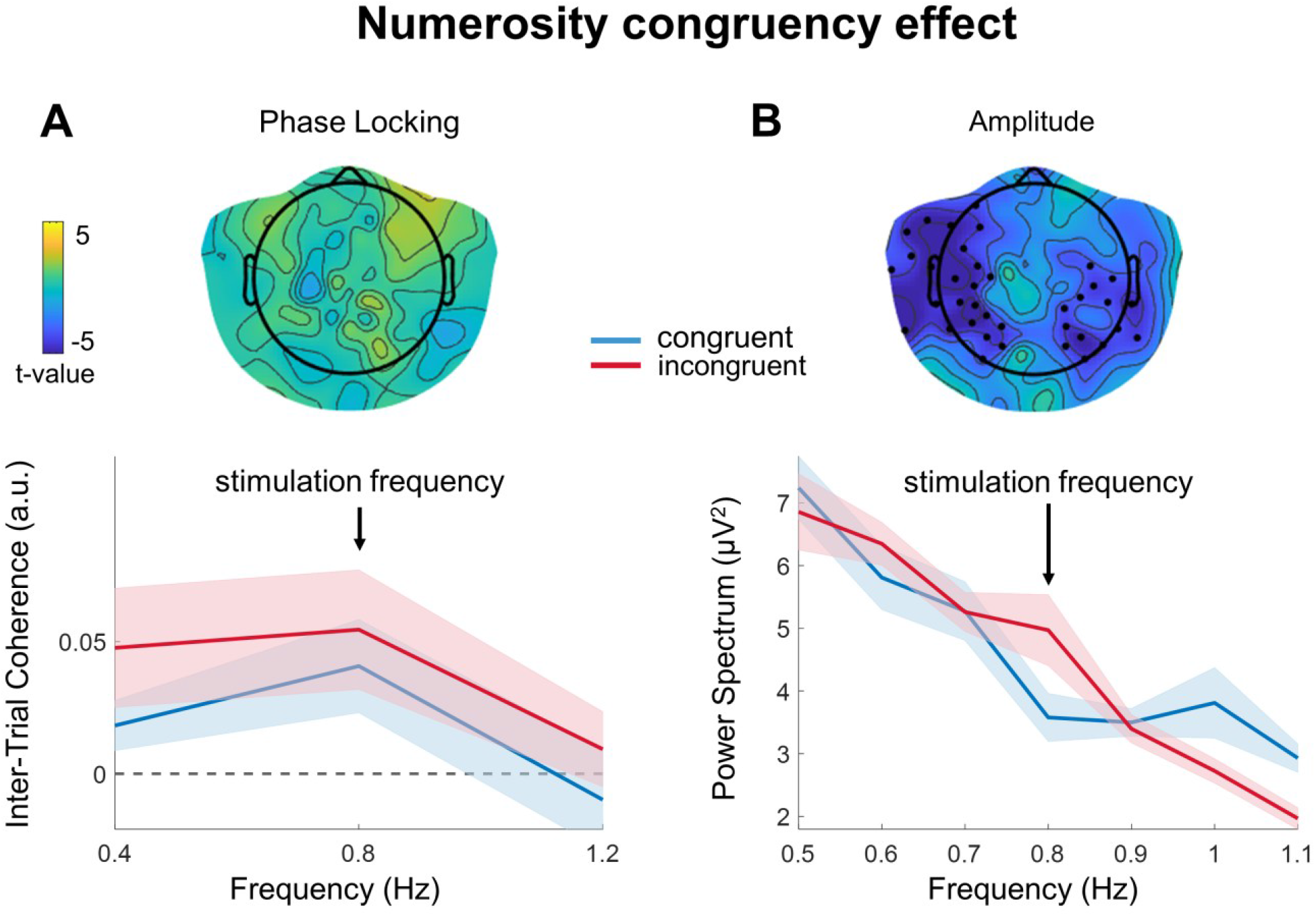
Neural adaptation to cross-modal numerosity congruency. (A) Top: Statistical map (t-value, paired t-test; black dots indicate statistically significant clusters) of the difference between the ITC in the congruent vs. incongruent condition. No significant difference is found. Bottom: For each condition, spectral profile of the difference between the ITC at the stimulation frequency (0.8 Hz) and the one computed from the null distribution, averaged over the significant cluster of the two conditions merged (Fig. 2A, top panel) (shaded contour indicates the SEM across subjects): for both conditions, a clear peak above the null-distribution level (marked by the dashed line at zero) emerges at the stimulation frequency, indicating that in both conditions the newborn brain temporally tracks the visual stimulation. (B) Top: Statistical map (as above) of the difference between the FTR in the congruent vs. incongruent condition. FTR is significantly lower in the congruent than in the incongruent condition in two bilaterally distributed clusters (left: *P*_*corr*_ < 0.001; right: *P*_*corr*_ < 0.014). Bottom: For each condition, mean of the single-subject difference between the PS and the background power averaged over the most significant (left) cluster (solid line) ± SEM (shaded contour), superposed on the average background power (dashed dark-gray line): while for the incongruent condition a clear peak emerges at the stimulation frequency, the PS for the congruent condition is flat, suggesting a cross-modal adaptation to numerosity.

Importantly, this effect was not influenced by the specific numerosity of the auditory sequences during familiarization: A between-subjects comparison of the difference of FTR values between congruent and incongruent conditions averaged over the most significant cluster (Fig. 3B) revealed no difference between newborns familiarized with 4 versus 12 syllables (between-subjects t-test: *P* = 0.84).

## Discussion

In this study, we investigated the existence of an abstract, supramodal code for number in the newborn brain by measuring the EEG responses of 0-3-day-old human newborns to visual arrays of items (4 or 12) that were either numerically congruent or incongruent to previously familiarised and concurrently presented auditory streams of a fixed number of syllables (4 or 12). Our results provide evidence that the newborn brain represents numerosity in an abstract, supramodal format, confirming with a direct neural measure previous behavioural evidence of cross-modal numerosity perception shortly after birth^3,4^. Specifically, in a large bilateral cluster of electrodes compatible with an associative parieto-temporal source^9^, we observed a significant decrease in the amplitude of the neural response to the visual stimuli that were numerically congruent with the previously familiarized and concurrently presented auditory ones, consistent with a cross-modal and cross-format repetition-suppression (adaptation) mechanism^7^. This effect could also be interpreted within a predictive coding framework, according to which adaptation effects reveal a reduction in prediction error for inputs that are expected^10^. This would suggest that newborns expect that concurrently presented visual and auditory stimuli have the same number of items, perhaps because they expect them to represent the same source. Although the current results do not allow us to disentangle these accounts, both require shared representations of numerosity across modalities.

Our paradigm was specifically designed to isolate numerosity from other magnitude-related dimensions. First, numerical information was conveyed through different sensory modalities, minimizing the possibility that congruency effects were driven by shared low-level sensory features. Second, following^3^, auditory sequences were equated across numerosities for extensive parameters (total duration), whereas visual arrays were controlled for intensive parameters (item size and density). As a result, numerosity was the only dimension systematically shared across the auditory and visual stimuli, making it the most parsimonious explanation for the observed cross-modal adaptation effect.

Our results are in line with previous EEG and fMRI-based repetition-suppression effects observed using visual numerosity stimuli in both 3-month-old infants^9^, preschool children^11^ and adults^12,13^, as well as with psychophysical findings of cross-modal numerosity adaptation between visual and auditory stimuli in adults^14^, and cross-modal decoding of numerosity in 3-month-old infants^5^ and adults^15^. At first glance, the reported cross-modal neural repetition-suppression effect for numerically congruent arrays might seem at odds with the behavioural findings that report longer looking times to such stimuli^3,4^ (but see^16^). However, repetition suppression reflects changes in neural encoding that do not necessarily imply a reduced motivational or attentional engagement^7^. Indeed, our data show that EEG phase locking to the visual stimuli was equivalent for congruent and incongruent stimuli, indicating comparable levels of attentional tracking across conditions.

Together, these findings support the view that numerosity constitutes a core dimension of human perception, supported by neural systems that are operational from the very first days of life. By providing evidence for an abstract neural code for number in newborns, this study contributes to a deeper understanding of the origins and functional properties of the early human number sense.

## Methods

The study was approved by the local ethical committee for clinical research (Comitato Etico per le Sperimentazioni Cliniche, Azienda Provinciale Servizi Sanitari, Province of Trento, Italy) and was performed in the maternity ward of Rovereto Hospital Santa Maria del Carmine. Parents were informed about the content and goal of the study and gave their written informed consent.

### Subjects

Twenty-one newborns were included (11 females; mean age 35 ± 13 hours, range 14 to 62 hours). All were healthy (APGAR(5 min) = 10), born full term (gestation age: 39.7 ± 1.0 weeks), and of normal birthweights (average weight 3.32 ± 0.28 Kg, range 2.70-3.75 Kg). Forty-four additional newborns participated but were excluded from the final sample either because they did not complete the study (criteria: attend both stimulus conditions for at least 20 s each) due to inattentiveness or fussiness (26), falling asleep (3), technical problems (3) or because their data contained too many EEG artifacts (mainly due to movements or high electrode impedance) (12).

### Stimuli

Stimuli were presented using the Psychtoolbox 3.0.12 for Windows in Matlab R2014a (Natick, MA). Infants were first familiarized with a continuous auditory stream, which consisted of sequences of syllables each repeated a fixed number of times (either 4 or 12). During familiarization, all the sequences had the same numerical value for any given infant; this number was randomly balanced across participants. During this phase newborns were placed in front of a black screen. Infants were then tested with visual images presenting either 4 or 12 items, i.e. the same number and a different number of items relative to auditory stimuli. Eight different syllables (*co, la, lo, ma, ni, pe, su, ti*) of variable duration and voice (4 male, 4 female) were recorded to construct the sequences. Sequences were equated in duration across numbers, hence each syllable was shorter for the larger number (12) than for the smaller number (4). The average duration of syllable sequences was 3.0 s. Successive sequences were separated by a variable silent interval of 1–3 s duration.

Test images were prepared with a variant of a program used in^3^. Each image presented a set of brightly colored, simple geometrical shapes. All the objects were identical within each image; four different shapes (squares, diamonds, triangles and circles) combined with four colors (red, green, yellow and purple, respectively) were used across the four test images. The order of appearance of the different shapes, and their pairing with the different numbers (congruent or incongruent, 4 or 12) were randomized across participants. Stimuli were presented dynamically with sinusoidal contrast modulation (0-100%) at a rate of 0.8 Hz (1 cycle = 1.25 s), unsynchronized with the onset of auditory sequences (Fig. 1). We used sinusoidal contrast modulation instead of squared on–off dynamics, both to minimize nonlinear effects in the brain frequency response^17^ and to make the stimulation more pleasant to the babies. Following^8^, the slow presentation rate (0.8 Hz) was chosen to ensure that newborns fully perceived the stimuli at each cycle of the periodic, peekaboo-like presentation. Stimuli were presented on a black circle (visual angle = 25°) superposed to a gray background.

In contrast to the auditory stream, in the test images the intensive parameters were equated across numbers (item area = 6.25 cm^2^ for all items, square/diamond side = 2.5 cm, circle diameter = 2.8 cm, triangle side = 3.8 cm), such that the extensive parameters (summed luminance, total surface occupied by the array) increased with number. As a consequence, if infants attended to intensive parameters and were able to generalize these parameters across modalities, they would respond equally to all of the test images; if infants attended to extensive parameters and showed cross-modal generalization of those parameters, they should exhibit the same test preference regardless of the familiarization conditions.

### EEG recordings

EEG was recorded with a high-density (125 electrodes) Geodesic EEG system (GES400 EGI, USA) referenced to the vertex. Scalp voltages were amplified and digitized at 250 Hz.

### Procedure

Newborns were tested in a calm, dimly illuminated space in the maternity leave, seated on the lap of a trained student in front of an LCD screen (60 × 33.8 cm; distance from eyes to screen: about 30 cm) slightly tilted towards the subject, while wearing the EEG cap. The student was instructed to maintain visual attention on the newborn. However, we cannot exclude that sometimes they also looked at the screen. Video recording from a hidden camera on the top of the screen ensured on-line monitoring of the infant. The newborn’s parents, when present, were off the sight of the infant (separated by a curtain), and instructed to keep silent during the recordings.

Each recording started with the familiarization phase: infants were exposed to the continuous auditory stream while the screen was blank. After 1.5 minutes, the visual stimulation started with a distractor consisting of a black and white spiral looming towards the center of the screen on a gray background. As soon as the newborn started to fixate the center of the screen, the periodic stimulation with test images started. Each trial was presented for 40 cycles (50 seconds) or until the subject stopped fixating, and it ended with a blank screen. Then the alternative condition was presented. In order to maximize the duration of the EEG recordings for each condition, this alternation continued until the newborn became fussy or asleep. The starting condition (congruent or incongruent) was balanced across newborns. Fixation times were computed off-line by two independent coders who reviewed the video recordings blindly with respect to the experimental conditions. Mean fixation time did not differ between conditions (paired t-test: *F*(1,20) = 1.30, *P* = 0.268; congruent: 110.0 ± 51.8 s; incongruent: 98.1 ± 59.1 s), indicating that the observed differences in EEG responses cannot be attributed to differences in data statistics. This absence of a fixation-time difference may reflect two modifications introduced relative to the original paradigm^3^ to maximize EEG recording duration for both conditions: (a) we set the maximum trial duration to 50 s (compared with 60 s in Izard et al.), and (b) we kept presenting trials until the infant became fussy or fell asleep, whereas Izard and colleagues systematically ended the experiment after collecting four trials per infant.

### EEG data analysis

Data analysis was performed with the EEGLAB toolbox^18^, the Fieldtrip toolbox^19^ and custom-made software based on MATLAB R2024a (Natick, MA).

### EEG pre-processing

EEG data were band-pass filtered (high-pass filter: EEGLAB function clean_drifts (transition band: 0.15-0.3 Hz); low-pass filter: default EEGLAB low-pass filter (40 Hz)) and segmented in blocks corresponding to fixation intervals. To optimize the trade-off between artifact removal and correction with the typically short and noisy newborn EEG data, we performed EEG artifact detection and correct/removal by combining two automatic pipelines in a fully automatic fashion. In the first step, we used NEAR^20^ – an artifact removal pipeline specifically designed for newborn EEG data – to remove heavily artefacted segments from the data. More specifically, we first removed bad channels (LOF parameter = 2.5, adapted to newborns) and then we identified the bad segments by applying ASR with a relatively high threshold (k=50). In a second step, we applied the bad segment rejection on the filtered data prior to bad channel correction, and ran APICE^21^ – an automatic artifact correction pipeline for developmental cognitive studies – to both correct and remove the residual artifacts. A final check by visual inspection of the data was performed to reject the (very few) residual artefacts. The automaticity of this process guarantees that data from all subjects and both experimental conditions are processed with the very same criteria, an important point given the shortness and noisiness of newborn EEG data. After artifact rejection, the duration of clean EEG data per condition was, on average, 135.7 s (congruent, 69.5 ± 35.5 s; incongruent, 66.1 ± 38.8 s), with no statistical difference among the two conditions (paired t-test: *P* = 0.54). The resulting signals were mathematically referenced to the average of the 125 channels.

### Power spectrum analysis

We computed the power spectrum analysis as in^8^. To obtain a high frequency resolution of the power spectrum with one bin centered on the stimulation frequency (0.8 Hz), we chose the longest possible epoch length compatible with the shortness of the recorded newborn EEG data^22^, i.e. 8 stimulation cycles (10 s), resulting in a frequency resolution of 0.1 Hz. EEG data from each block were segmented in partially overlapping epochs of 10 s (overlap varied between ½ and 3/4 of epoch length to include all time points). For each electrode, the Fourier transform *F*(*f*) of each epoch was calculated using a fast Fourier transform algorithm (Matlab function FFT). The power spectrum was calculated from these Fourier coefficients as the average over epochs of the single-epoch power spectrum: *PS*(*f*) = ⟨*F*(*f*) × *F*\*(*f*)⟩_*ep*_. The Frequency-Tagged Response (FTR) at the tag frequency (0.8 Hz) was calculated as the ratio between the power spectrum at the tagged frequency and the value at 0.8 Hz of the power-law fit of the power spectrum estimated from the 6 neighboring frequency bins (+-0.3 Hz), where the power-law fit was computed by fitting a line to the logarithm of the power at the 6 neighboring frequency bins (Matlab function Polyfit).

### Phase locking

To compute phase locking of the EEG signal to the phase of the oscillatory stimulation (also called inter-trial phase coherence), we chose a shorter epoch length (2 stimulation cycles = 2.5 s, resulting in frequency bins of 0.4 Hz), as it provides a better sampling of the phase distribution for short data length, a necessary condition for a reliable estimate of phase locking^22^. Consecutive non-overlapping epochs time-locked to the onset (contrast=0) of the stimulus were extracted from the EEG data blocks.

We estimated phase locking by using the Inter-Trial Coherence (ITC^23^):

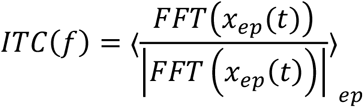

### Statistical analysis

We tested the statistical significance of all effects with the non-parametric cluster-based test^24^ implemented in Fieldtrip^19^. This method allows statistical testing with no need of a priori selection of spatial ROIs because it controls for multiple comparisons by clustering neighboring channel pairs that exhibit statistically significant effects (test used at each channel point: dependent-samples *t* statistics, threshold: *p*=0.05, one-sided for the overall visual entrainment effect, two-sided for the difference between congruent and incongruent conditions) and using a permutation test to evaluate the statistical significance at the cluster level (Montecarlo method, 3000 permutations for each test). The *p* value of each statistically significant cluster is indicated as *P*_*corr*_ to mark that it is “corrected” for multiple comparisons. Specifically, the channel-level tests for each measure were the following.

Power spectrum/FTR: We evaluated the statistical significance of the entrainment to the oscillatory visual stimulation by comparing the logarithm of the power at the tag frequency with the logarithm of the background power estimated by the power-law fit described above. Differences between conditions were evaluated by comparing the logarithm of the relative FTRs.

ITC: We evaluated the statistical significance of the ITC at the tag frequency by comparing it with the ITC of surrogate data generated under the null hypothesis of non-entrained oscillatory activity: we generated 100000 random phase vectors of the same length of the original phase data (to account for finite-size effects), we computed the ITC for each vector and we took the average ITC as a stable estimate of the surrogate ITC null distribution.

The effect size was estimated with a post-hoc analysis on the statistically significant cluster by computing Cohen’s *d* as the ratio between the mean and the standard deviation (across subjects) of the effect averaged over all the electrodes of the cluster.

## Data availability

The anonymized EEG datasets and custom analysis scripts are publicly available at https://osf.io/8xzeq/overview?view_only=9e6d4eb99c1348be9e70978be8fbfb30

## Acknowledgements

We thank the staff of the Pediatrics and Obstetrics/Gynecology Units of Rovereto Hospital, the newborns involved in the study and their parents, and Marta Mosele and Giulia Vigna for invaluable assistance with data collection.

## Author contributions

Conceptualization: M.B., V.I. and M.P.; Investigation: M.B., E.E. and I.P.; Methodology: M.B.; Formal analysis: M.B. and E.E.; Resources: U.P. and F.T.; Funding Acquisition: G.V. and M.P.; Writing – Original Draft Preparation: M.B., E.E. and M.P.; Visualization: M.B and M.P.; Writing – Review & Editing: all authors.

